# Bayogenin 3-O-Cellobioside is a novel non-cultivar specific anti-blast metabolite produced in rice in response to *Pyricularia oryzae* infection

**DOI:** 10.1101/647636

**Authors:** Justice Norvienyeku, Lili Lin, Abdul Waheed, Xiaomin Chen, Jiandong Bao, Ammarah Shabbir, Lianyu Lin, Zhenhui Zhong, Wajjiha Batool, Sami Rukaiya Aliyu, Jie Zhou, Guodong Lu, Zonghua Wang

## Abstract

Rice cultivars from *japonica* and *indica* lineage possess differential resistance against blast fungus on an account genetic divergence. Whether different rice cultivars also show distinct metabolomic changes in response to *P. oryzae*, and their role in host resistance, are poorly understood. Here, we examine the responses of six different rice cultivars from *japonica* and *indica* lineage challenged with *P. oryzae*. Both susceptible and resistant rice cultivars expressed several metabolites exclusively during *P. oryzae* infection, including the saponin Bayogenin 3-O-cellobioside. Bayogenin 3-O-cellobioside level in infected rice directly correlated with their resistant attributes. These findings reveal, for the first time to our knowledge that besides oat, other grass plants including rice produces protective saponins. Our study provides insight into the role of pathogen-mediated metabolomics-reprogramming in host immunity. The correlation between Bayogenin 3-O-Cellobioside levels and blast resistance suggests that engineering saponin expression in cereal crops represents an attractive and sustainable disease control strategy.

## Introduction

Rice blast disease accounts for yield losses that could feed about 60 million rice consumers annually(1). *Pyricularia oryzae* (Syn: *Magnaporthe oryzae*), the fungus that causes rice blast disease(2), is ubiquitously present in all rice-producing regions across the globe. In addition to rice, blast fungus can infect wheat, barley, millet, sorghum, rye, and other cultivated as well as non-cultivated grass plants, and is therefore considered one of the most important plant pathogens (3).

Rice blast fungus initiates infection by producing asexual spores (conidia) which disperse and attach onto healthy host tissues (4). In favorable conditions, the spore propagule (inoculum) germinates and produces a short hyphae-like structure typically from the apical cell called germ tube which later differentiates into a bulbous infectious structure known as appressoria (5). The appressoria further differentiates into rigid and robust penetration structure (penetration-peg) which is engaged by the blast pathogen to physically rupture the cuticle of susceptible host plants for successful invasion, colonization of host cells and the manifestation of blast symptoms (6, 7).

Current rice blast control strategies rely heavily on the use of rice cultivars with inherent basal resistance against the pathogen, as well as the breeding of resistant (R)-gene aided cultivars including CO39, Pi-b, Pi-4b, Pi-a, Pi-9, Piz-t, Pi-pita, Pi-gm, Pi9, Pi2, among others (8-10). However, rice blast evolves rapidly to overcome R-gene supported resistance (11), and inherent basal resistance does not offer full immunity against blast fungus. Rice cultivars from *japonica* and *indica* sub-species possess differential resistance against blast fungus, with *japonica* rice mainly pose inherent resistance whereas *indica* rice poses R-gene mediated resistance (9). Although rice-specific secondary metabolites (phytochemicals) such as oryzalexins, phytocassanes, momilactone, and sakuranetin can protect rice from bacterial and fungal pathogens, including blast fungus (12, 13), these compounds have not been extensively exploited as potent defense molecules against blast pathogen, largely because most of these metabolites are cultivar-specific and were identified enhance in response to phytohormone treatment rather than pathogen treatment. Current advances in metabolomics technologies enable the identification of phytochemicals and defense signaling molecules that are induced in plant tissues during host-pathogen interactions. Here, we investigated how *P. oryzae-*mediated metabolome reprogramming affects the susceptibility or resistance of different rice cultivars to *P. oryzae*.

## Results

### Raw leaf extracts from inoculated rice seedlings inhibit infectious development of *P. oryzae*

Firstly, we confirmed the susceptibility and resistance of six different rice cultivars (CO39, LTH, NPB Pi-B, Pi-4B, and Pi-gm) from both *indica* and *japonica* lineage (supplementary table 1) to *P. oryzae*. Briefly, we spray-inoculated three-week-old seedlings with conidia suspensions prepared from the *P. oryzae* Guy11 strain, incubated them in a dark and humid chamber, and transferred them to a growth chamber. At 7-days post inoculation, we assessed the type and severity of lesions on leaf tissues according to a published rice blast lesion scoring index (14). We found that the CO39, NPB, and LTH cultivars were highly susceptible, with a higher number of severe blast lesions (type 4 and 5 lesions), whereas Pi-gm, Pi-4b, and Pi-b cultivars displayed moderate to complete immunity against blast fungus (Figure.1B).

**Figure 1:**
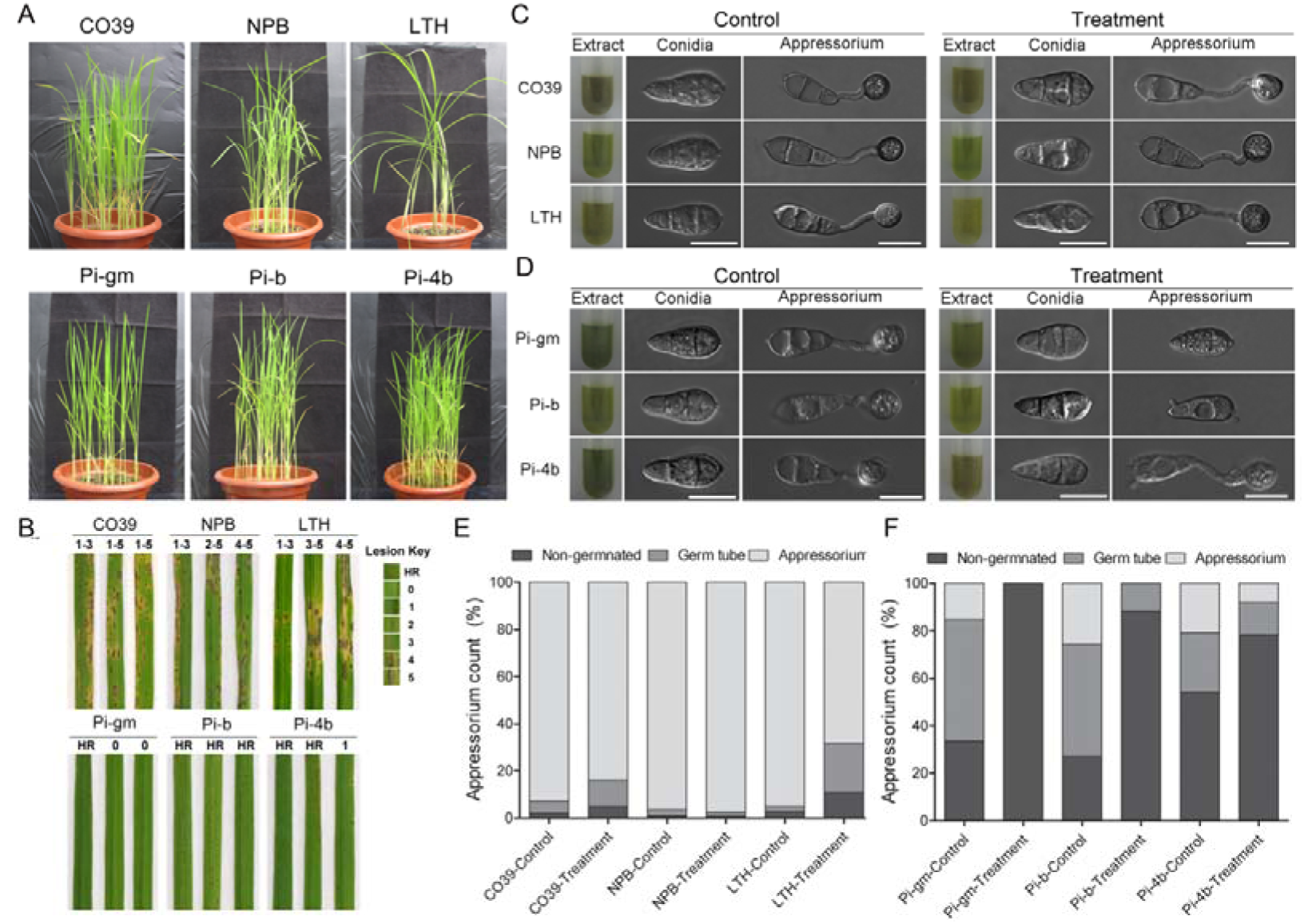
Crude leaf extracts from pre-inoculated resistant rice cultivars significantly inhibits germination and appressorium formation in *P. oryzae*. **(A)**. Showed blast symptoms and the susceptibility index of homogenously susceptible rice cultivars sprayed inoculation with conidia suspensions of *P. oryzae*. **(B)** Showed the development blast symptoms and resistance attributes displayed by moderate to completely resistant rice cultivars sprayed inoculation with conidia suspensions of *P. oryzae*. Note hypersentive response (HR)-0 represent complete resistance, lesion type 1-2 represent moderate resistance while lesion type 3-5 signifies complete susceptibility.**(C)** Exhibit germination and appressorium formation characteristics of rice blast fungus spores treated with total crude extracts obtained from pre-inoculated and non-inoculated susceptible rice cultivars. **(D)**. The micrograph display the inhibitory effects of total crude extracts obtained from pre-inoculated and non-inoculated resistant rice cultivars on the germination of rice blast fungus spores. **(E)**. The stacked column graph is a statistical presentation of *P. oryzae* spores treated with total crude extracts from susceptible rice cultivars. **(F)**. The stacked column graph is a statistical presentation of *P. oryzae* spores treated with total crude extracts from resistant rice cultivars. Results for infection assay (**A, B)** was obtained from three biological experiments with five technical replicates n=600 with a conidia concentration of 3.0×10*5. Microscopy examination and statistical analysis, **C, D, E, & F** (n=750), Scale bars, 10 μm.

We hypothesized that metabolites produced after inoculation with rice blast might inhibit infectious development of *P. oryzae*. To test this hypothesis, we extracted crude extracts from inoculated rice seedlings, as well as from non-inoculated controls. We used the crude leaf extracts to wash conidia from the *P. oryzae* Guy11 strain, prepared conidia suspensions, and inoculated a hydrophobic coverslip to induce appressorium formation *in vitro*. Crude extract from the untreated resistant cultivars inhibited conidia germination and appressorium formation, whereas crude extracts from the untreated susceptible cultivars had no inhibition effect (Figure. 1C-D). However, in 5/6 cases, the crude leaf extracts from the inoculated seedlings inhibited germination and appressorium formation more than crude leaf extracts from the corresponding non-inoculated control seedlings (Figure. 1E-F). Thus, indicate the likely presence of anti-blast metabolites in crude extracts from both susceptible and resistant cultivars show increased production of upon inoculation *P. oryzae*.

### Resistant rice cultivars accumulate higher levels of Bayogenin 3-O-cellobioside upon inoculation with *P. oryzae*

To monitor the changes in rice seedlings due to *P. oryzae* infection, we performed metabolomic analysis of inoculated and non-inoculated rice cultivars. We spray-inoculated two-week-old susceptible (CO39, NPB, and LTH) and resistant (Pi-gm, Pi-4B, and Pi-B) rice seedlings with conidia suspensions along with non-inoculated controls (Figure.2A), harvested leaf tissues at 12-hours post inoculation, and extracted metabolites in methanol using QTOF-UPHPLC (see Methods). Also to ensure exclusion of fungi specific metabolites, we analyzed the metabolomes of *P. oryzae* at different developmental stages including vegetative growth (mycelium stage), conidia (aberrant conidia/ resting stage) and conidia germination and appressorium formation stage (infectious development stage) (see Methods, and (15). Principal Component Analysis (PCA) of the data revealed the reproducible identification of metabolites in at least five out of the seven independent repeats (Figure.2A).

**Figure 2:**
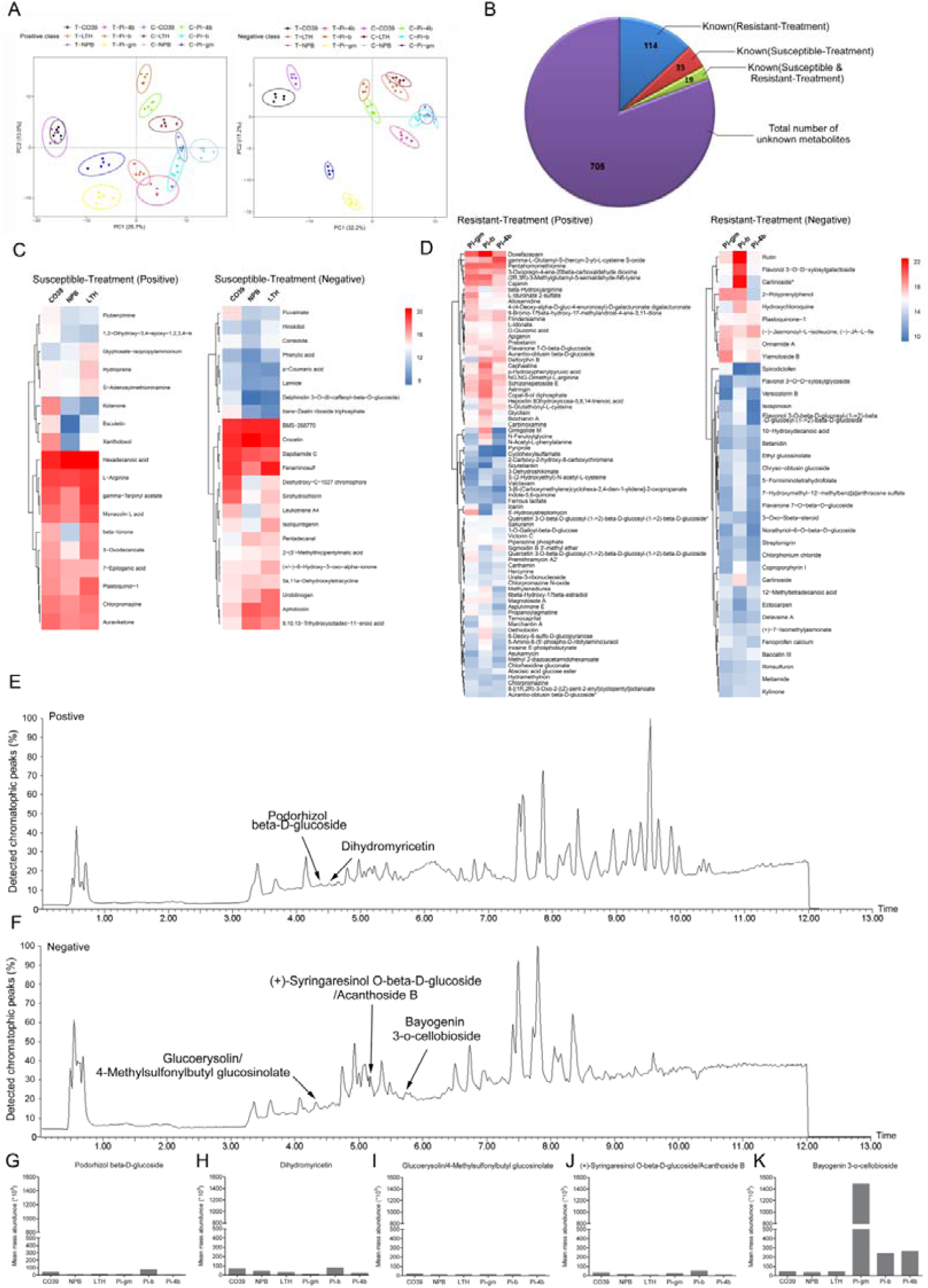
Susceptible and resistant rice cultivars undergo common and differential metabolome reprogramming in response to *P. oryzae* infection. **(A).** PCA showed the number of common and differential metabolites recorded exclusively in susceptible and resistant rice cultivars challenged with rice blast fungus under positive (Pos.+) ionization mode. **(B)**. Showed the number of common and different metabolites recorded in susceptible and resistant rice cultivars inoculated with the rice blast fungus under both negative (Neg-.) and Pos.+) ionization mode. **(C)**. The graphical matrix showed the list, clusters and abundance intensity of metabolite generated exclusively in susceptible rice cultivars in response to rice blast fungus infection under (Neg.-) ionization mode. **(D)**. The heat-map showed names and cluster of unique metabolites identified exclusively in resistant rice cultivars inoculated with *P. oryzae* under (Pos.+) ionization mode along with their comparative mass abundance intensities across the respective cultivars. **(E)** Represent the chromatograph of metabolites identified exclusively in both susceptible and resistant rice cultivars inoculated with the rice blast fungus under positive (Pos.+) ionization mode. **(F)** Displayed the chromatograph of common metabolites identified in susceptible and resistant rice cultivars challenged with the rice blast fungus under negative (Neg.-) ionization mode. **(G-K).** Displayed mass baundances recorded for non-cultivars specific metabolites identified in both susceptible and resistant rice cultivars under under positive (Pos+) and negative (Neg-.-) ionization mode. Structures for the respective metabolites are presented in supplementary figure 3. Metabolomics data was obtained from one one biological replicate with six technical replicates (n= 240, seedling population per replicate =40). Metabolites with a a mass abundance ≥ 1000, mass error ≤±3 T-test P-value (q-value) □0.05, relative standard deviation (RSD) □30% est P-value (q-value) □0.05, relative standard deviation (RSD) □30% and a minimum score of 25% were filtered and used for comparative metabolomics analysis **(A, B, C, & D).** Quantitative inter-metabolomics variation between groups was analyzed with ANOVA **(G, H, I, J, &K).**

To uncover disease-relevant metabolites from the whole-metabolome profiles, we developed a robust filtering system to diminish unwanted biological variables, background noise, and biologically insignificant metabolites (see Supplementary Fig.1a, Methods, and (15). We uncovered ∼2121 metabolites in susceptible rice seedlings, and ∼3450 metabolites in resistant rice seedlings (Supplementary Table 2a). Kyoto Encyclopedia of Genes and Genomes (KEGG) compound enrichment analysis (16) revealed that 883 metabolites were produced exclusively in infected seedlings (Figure.2 A &B) and (Supplementary Figure.1b). Of these 883 metabolites, 705 were unknown. Of the 178 metabolites with a KEGG code, 18 were present in both susceptible and resistant rice cultivars after inoculation, 114 were present exclusively in the susceptible cultivars after inoculation, and 45 were solely present in the resistant cultivars after inoculation (Figure.2B-D).

To determine the function of the 18 metabolites present in both susceptible and resistant rice seedlings upon infection, we used their compound codes, chemical formulae and common names to search publicly available metabolite libraries, including KEGG compound, KEGG BRITE, PubChem Compound, The Human Metabolome Database, The Small Molecule Pathway Database, and The Toxin and Toxin Target Database (17). We found that 5/18 metabolites are known to be functional phytochemicals: Podorhizol beta-D-glucoside, Bayogenin 3-O-cellobioside, (+)-Syringaresinol O-beta-D-glucoside, 4-Methylsulfonylbutyl glucosinolate, and Dihydromyricetin (Figure.2E-F) and (Supplementary Figure. 3). Indeed, Podorhizol beta-D-glucoside and 4-Methylsulfonylbutyl glucosinolate are known to enhance plant immunity against diverse pathogens (18, 19).

**Figure 3:**
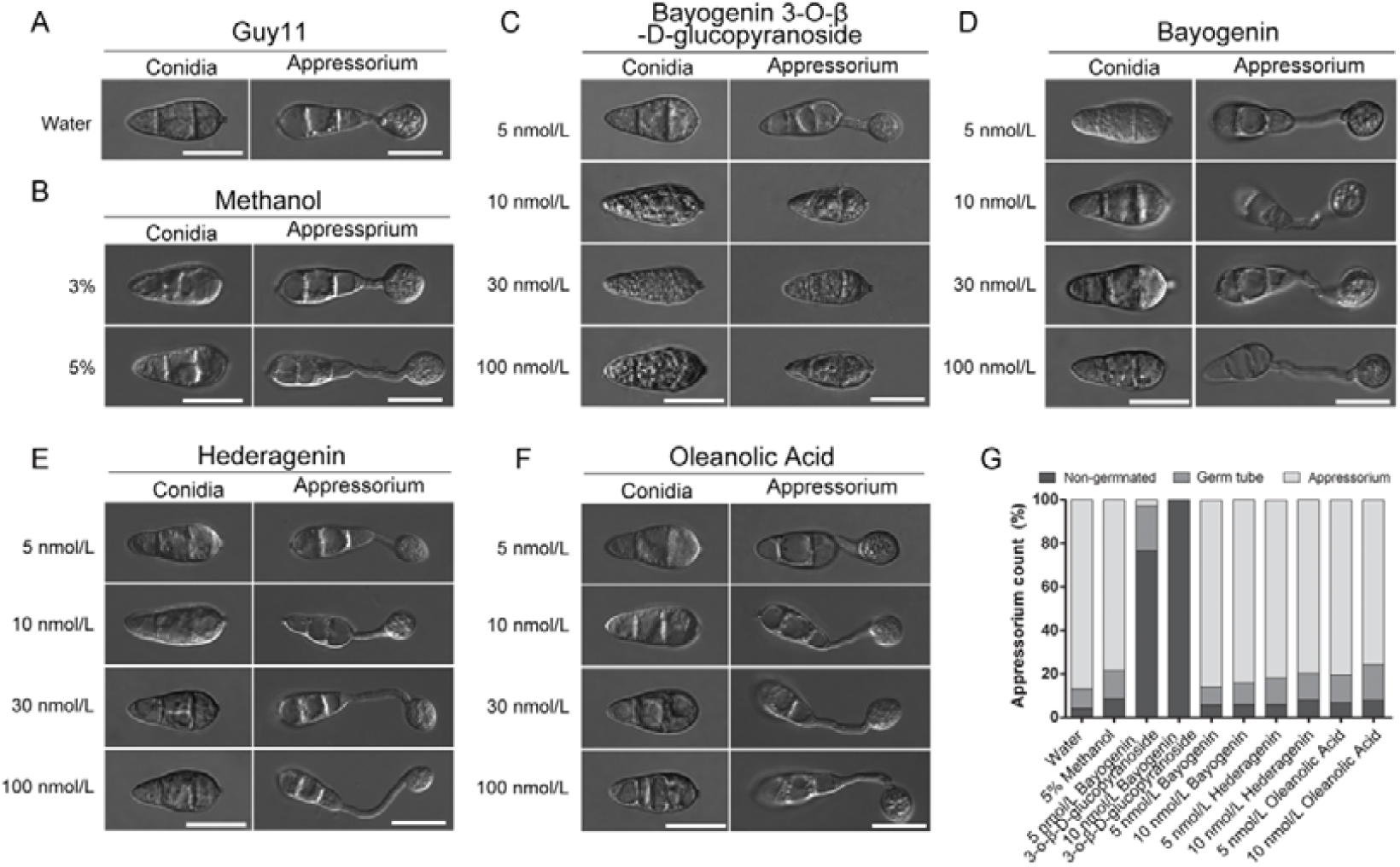
Glycosylated Bayogenin exclusively inhibits the germination and appressorium formation in conidium produced by the rice blast fungus. **(A)**. Represents the negative control group that portrays the morphological characteristics of appressorium produced by *P. oryzae* conidia suspensions prepared with double deionized water (ddH2O). **(B).** Displayed the impact of 3% and 5% percent methanol on conidia germination and appressorium formation (positive control group). **(C).** The micrograph represents the inhibitory effects of different concentrations of glycosylated Bayogenin (Bayogenin 3-O-β-D-glucopyranoside) on conidia germination and appressorium development. **(D)** Showed the influence of non-glycosylated Bayogenin on germ tube speciation and appressorium development in *P. oryzae*. **(E)** Showed the impact of Hedegeranin on conidia germination, and appressorium formation in *P. oryzae* conidium. **(F)**. Showed germination and appressorium formation characteristics of *P. oryzae* conidia treated with Oleanolic acid. **(G).** The stacked column bar graph is a statistical display of the effect of different saponins on conidia germination and appressorium morphogenesis. Statistical computation was performed using average values obtained from three biological experiments with three replicate each time for all treatment (n=750). Scale bars, 10 μm.

Intriguingly, unlike the other four metabolites, the levels of Bayogenin 3-O-cellobioside were relatively low in the inoculated, susceptible rice cultivars compared to in the inoculated, resistant group (Figure. 2 G-K). Furthermore, we observed that Bayogenin 3-O-cellobioside levels were about ∼1000-fold higher in the completely resistant cultivar Pi-gm than in the moderately resistant cultivars Pi-4b and Pi-b (Figure.2K). Thus, Bayogenin 3-O-cellobioside levels increase specifically upon inoculation of rice seedlings with *P. oryzae* and directly correlate with the extent of resistance. Our results suggest that Bayogenin 3-O-cellobioside is a novel general defense molecule produced in response to rice blast fungus infection, in both susceptible and resistant rice cultivars.

### Glycosylated Bayogenin inhibits infectious development of *P. oryzae in vitro*

Saponins are glycosylated triterpenoids, generated in many plants as a secondary metabolite. Different saponins display potent insecticidal and fungicidal activities against a broad range of plant parasites (20, 21). Unlike dicotyledonous plants, monocots such as rice are considered triterpenoid-poor plants (22). Avenacin is a saponin in oat (Avena spp) that provides defense against soil-borne fungal pathogens. Bayogenin 3-O-cellobioside is a glycosylated saponin, consisting of a non-sugar aglycone (Bayogenin) linked to a sugar glycone (Cellobioside). Because Bayogenin 3-O-cellobioside is not commercially available, we tested both non-glycosylated Bayogenin and glycosylated Bayogenin (Bayogenin 3-O-β-D-glucopyranoside) in an *in vitro* conidia germination and appressorium formation assays. We prepared titrations of these compounds and used them to wash conidia produced by the *P oryzae* Guy11 strain. Subsequently, we prepared conidia suspensions and inoculated a hydrophobic coverslip. We observed a dose-dependent inhibition of conidia germination and appressorium formation *in vitro* with increasing concentrations of Bayogenin 3-O-β-D-glucopyranoside (5 nM/L-100 nM/L) (Figure.3 C and G). In contrast, increasing the concentrations of the non-glycosylated Bayogenin did not affect germination or appressorium formation, nor did the control treatments (Figure.3 D and G D).

To ascertain whether other saponins would similarly inhibit conidia germination and appressorium formation, we used the *in vitro* assays to test two closely related saponins, Hedegeranin, and Oleanolic acid. Despite their reported insecticidal effects (23, 24), we found that washing conidia with 5-100 nM/L of Hedegeranin or Oleanolic acid did not adversely affect conidia germination and appressorium morphogenesis of *P oryzae in vitro* (Figure.3 E, F, and G). Together, these findings suggest that glycosylated Bayogenin, such as the Bayogenin 3-O-cellobioside, and Bayogenin 3-O-β-D-glucopyranoside are potent phytochemicals that specifically inhibit infectious development of *P oryzae*.

### Differential expression of steroid biosynthesis enzymes during rice-*P. oryzae* interaction

Genes encoding enzymes involved in steroid biosynthesis, including β-amyrin synthases, uridine diphosphate (UDP) glucuronyltransferases/ glycosyltransferases (UDP/GTs), appear to mediate the biosynthesis of saponins in plants (22, 25, 26). We identified a total of 6 rice β-amyrin synthases and 145 putative rice-specific UDP-GTs from the publicly available glycosyltransferase database (27). Our bioinformatics analysis revealed that the rice UDP-GTs family could be classified into 33 sub-families (groups) based on the alignment of a shared motif (Figure.4A). Further, 75 of this putative rice UDP-GTs are within genes clusters on multiple chromosomes (Figure. 4B).

**Figure 4:**
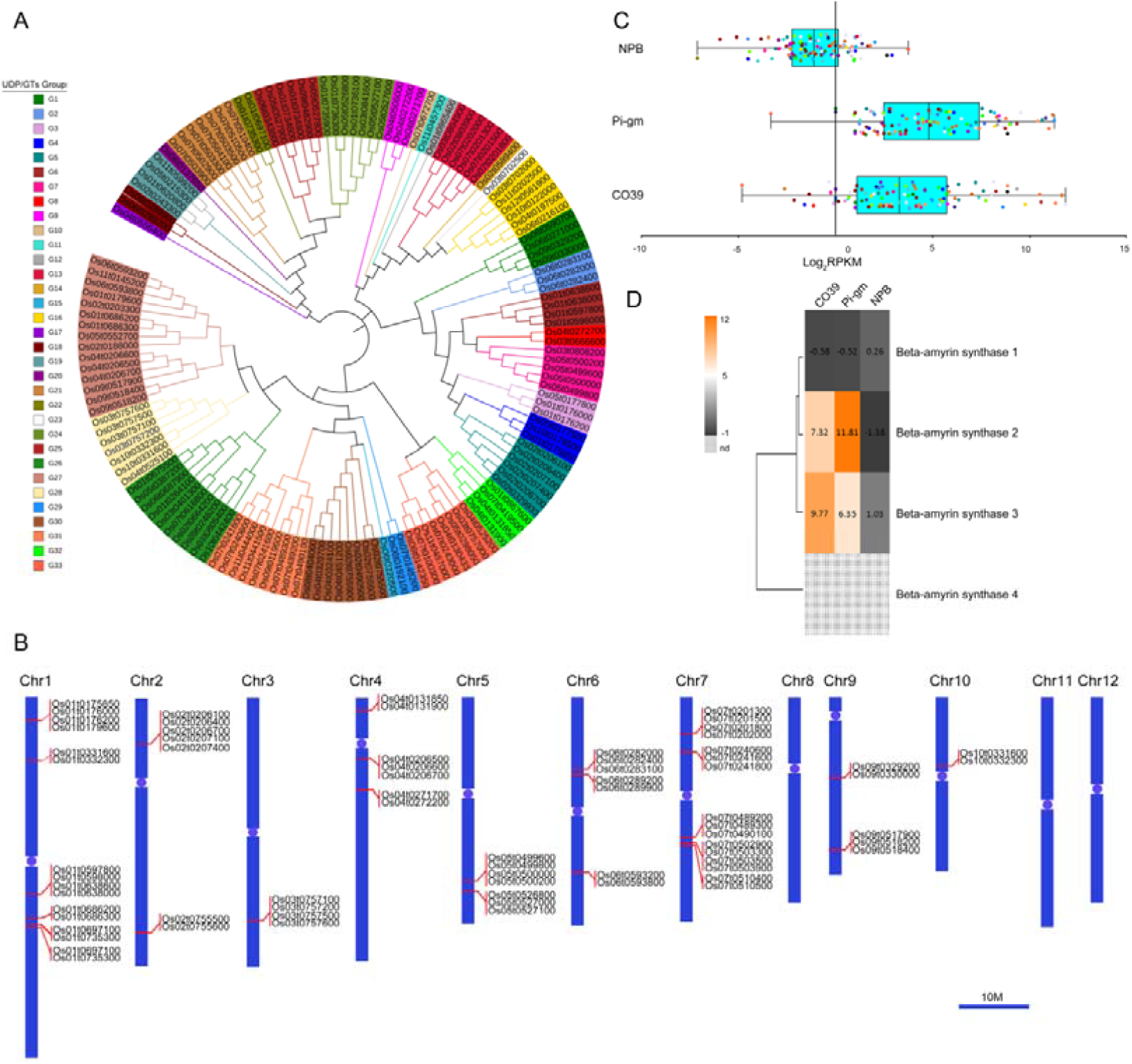
Clustering and expression profile of putative saponin biosynthesis gene in rice during *P. oryzae* infection. **(A)**. the Neighbor-joining tree showed the clustering 145 rice diphosphate glucuronyltransferases (UDP/GTs) into 33 different sub-families (groups) based on conserved alignable motifs. Each group is defined by a colour shade and consists of 1-15 genes. **(B)**. Displayed the chromosomal distribution and locations of UDP/GTs gene clusters in rice. The blue each vertical bar with upper and lower or (long and short) arms represent rice chromosome. The position of the blue circle (connecting the upper and lower arms) on each chromosome indicate represent the centromeric region. The numbering (Chr1-Chr12) on top of each vertical blue bar corresponds to Chromosome number. Set of genes aligned to red solid lines projecting from the chromosome represent a single cluster, therefore the number of red solid lines on each chromosome reflects the number of UDP/GTs gene clusters identified on the respective chromosome. **(C)**. Showed the comparative expression pattern of 108 differentially expressed UDP/GTs susceptible (CO39 and NPB) and resistant (Pi-gm) rice cultivars challenged with the rice blast fungus. Note the expression these 106 was exclusively in response to infection and were not detected in the non-inoculated control groups. Each coloured dot in or outside box plot represent the unique expression level (log2 FPKM) of UDP/GTs gene in the treated rice cultivars with a P=≤0.05. The asterisks at the whiskers indicate the lower and upper outliers.

We hypothesized that increased expression of either β-amyrin synthases or UDP-GTs might support the increased production of Bayogenin 3-O-cellobioside in rice seedlings inoculated with *P. oryzae*. To examine how inoculation affects the expression of these genes in different cultivars, we analyzed resistant (Pi-gm) and susceptible (CO39, NPB) seedlings by RNA sequencing. We found that of the 102 UDP-GTs were expressed exclusively upon inoculation of Pi-gm, whereas 83 and 13 UDP-GTs were expressed exclusively upon inoculation of CO39 and NPB, respectively (Fig.7C) and (see Supplementary Table 3). Also, two β-amyrin synthase genes were significantly (∼5-12 fold) up-regulated in Pi-gm and CO39 rice cultivars in response to *P oryzae* challenge (Figure.4D). Thus, various UDP-GT and β-amyrin synthase genes are expressed only upon inoculation of rice with *P oryzae*, in both susceptible and resistant seedlings. Our findings also underscore the importance of β-amyrin synthases and UDP-GTs in enforcing host immunity (28, 29). This clustering, and the observation that a higher number of UDP-GT genes are expressed upon inoculation of resistant cultivars compared to susceptible cultivars suggesting that multiple UDP-GTs enhance rice immunity against blast fungus by producing diverse glycosides, likely including Bayogenin 3-O-cellobioside. Disruption of a β-amyrin synthase and UDP-GTs blocks avenacin biosynthesis in oat (30, 31). However, the extent to which β-amyrin synthases and UDP-GTs influence Bayogenin 3-O-cellobioside biosynthesis, as well as the resistance or susceptibility of different rice cultivars to *P oryzae*, remains to be determined.

## DISCUSSION

We report that the rice metabolite Bayogenin 3-O-cellobioside (saponin) is a novel, general defense molecule that accumulates exclusively in response to *P. oryzae* infection, in susceptible and resistant rice cultivars from both japonica and indica linages. Further, the levels of Bayogenin 3-O-cellobioside after inoculation with *P. oryzae* directly correlate with the resistance of the rice cultivars. Therefore, susceptible and resistant rice cultivars display metabolomic differences not only before infection but also after infection, which contribute to differential resistance to rice blast pathogen.

Saponins are glycosylated triterpenoids, generated in many plants as a secondary metabolite. Different saponins display potent insecticidal and fungicidal activities against a broad range of plant parasites (20, 21, 32). Unlike dicotyledonous plants, monocots such as rice are considered triterpenoid-poor plants (22). Avenacin is a saponin in oat (*Avena* spp.) that provides defense against soil-borne fungal pathogens (33). To our knowledge, Bayogenin 3-O-cellobioside represents the first example of a saponin produced in rice.

β-amyrin synthases and UDP-GTs are key enzymes involved in the biosynthesis of saponins in plants (34-36). We found that two out of four putative β-amyrin synthases genes and a total of 106 putative UDP-GTs were specifically expressed during rice blast fungus infection of different cultivars. These data suggest that rice, and likely other monocots, are genetically capable of generating saponins and other glycosylated steroids under defined conditions.

Our findings also underscore the importance of β-amyrin synthases and UDP-GTs in enforcing host immunity (28, 29, 37). The clustering of rice UDP-GTs at single loci on a limited number of chromosomes suggests that clusters are likely controlled by a common regulator (38, 39). This clustering, and the observation that a higher number of UDP-GT genes are expressed upon inoculation of resistant cultivars compared to susceptible cultivars, suggests that multiple UDP-GTs enhance rice immunity against blast fungus by producing diverse glycosides, likely including Bayogenin 3-O-cellobioside. Disruption of a β-amyrin synthase and UDP-GTs blocks avenacin biosynthesis in oat (30, 31). The extent to which β-amyrin synthases and UDP-GTs influence Bayogenin 3-O-cellobioside biosynthesis, as well as the resistance or susceptibility of different rice cultivars to *P. oryzae*, remains to be determined.

Bayogenin 3-O-cellobioside is a glycoside consisting of a non-sugar aglycone (Bayogenin) linked to a sugar glycone (cellobioside) (40). Glycosylation is required to transform saponins to their bioactive state (41, 42). We found that spores from rice blast fungus treated with glycosylated Bayogenin (Bayogenin 3-O-β-D-glucopyranoside) failed to germinate whereas spores treated with non-glycosylated Bayogenin germinated and progressed to form functional appressorium. The aglycone component of insecticidal saponins is not sufficient to prevent *Phyllotreta nemorum* from feeding on the tissues of susceptible P-type of *Barbarea vulgaris* (43). Similarly, glycosylation plays a crucial role in promoting the fungicidal activities of Bayogenin.

Glycosides contribute to plant resistance against a broad range of parasitic insects and herbivores (44). However, the glycosides Hederagenin, and Oleanolic acid, did not inhibit spore germination or development of rice blast fungus. Bayogenin 3-O-β-D-glucopyranoside, on the other hand, significantly inhibited the germination of *P. oryzae* spores, but did not adversely affect vegetative development of rice blast fungus (data not shown), though there is limited knowledge on the evolution of saponin biosynthesis in different plant families (45). Differences in saponin bioactivity could be due to the composition of the core structure, functional groups, and the affinity with which these amphiphilic compounds integrate and disrupt the integrity of the targeted biological membrane systems (32, 45). Glycosylated Bayogenin (specifically Bayogenin 3-O-cellobioside) appears to be a novel and potent anti-fungal metabolite generated in both susceptible and resistant rice cultivars, providing a chemical defense against rice blast fungus. Soyasapogenol glycosides (*Lupinus angustifolius* L) and avenacin A (*Avena strigose)* display specific antifungal activity against *Candida albicans* and *Gaeumannomyces graminis* var. *tritici*, respectively, without a corresponding insecticidal effect(41, 46).

## Conclusion

Inherent metabolite differences between distinct rice cultivars have been associated with their distinct morphological and physiological characteristics (47-49). However, little is known about pathogen-induced metabolite differences between various rice cultivars and the potential impact on resistance or susceptibility traits. Beyond Bayogenin 3-O-Cellobioside, we found that other previously reported defense-related metabolites, such as abscisic acid glucoside ester (50), aurantio-obtusin-β-D-glucoside(51), carlinoside (52, 53) and sakuranin (derivative of sakuranetin) (54-57), were specifically produced in resistant rice cultivars challenged with rice blast fungus. Thus, resistant rice cultivars possess a metabolomic advantage over susceptible rice cultivars both before and during infection.

Overall, we report for the first time that diverse cultivars of rice produce a novel saponin (Bayogenin 3-O-Cellobioside) with anti-blast properties upon rice blast infection. We propose that β-amyrin synthases and/or UDP-GTs support saponin biosynthesis in rice (Fig.5). Our study provides insight into pathogen-mediated metabolomic reprogramming in host plants, and its impact on the resistant or susceptibility. The correlation between Bayogenin 3-O-Cellobioside levels and blast resistance suggests that engineering saponin expression in cereal crops represents an attractive and sustainable disease control strategy.

**Figure 5:**
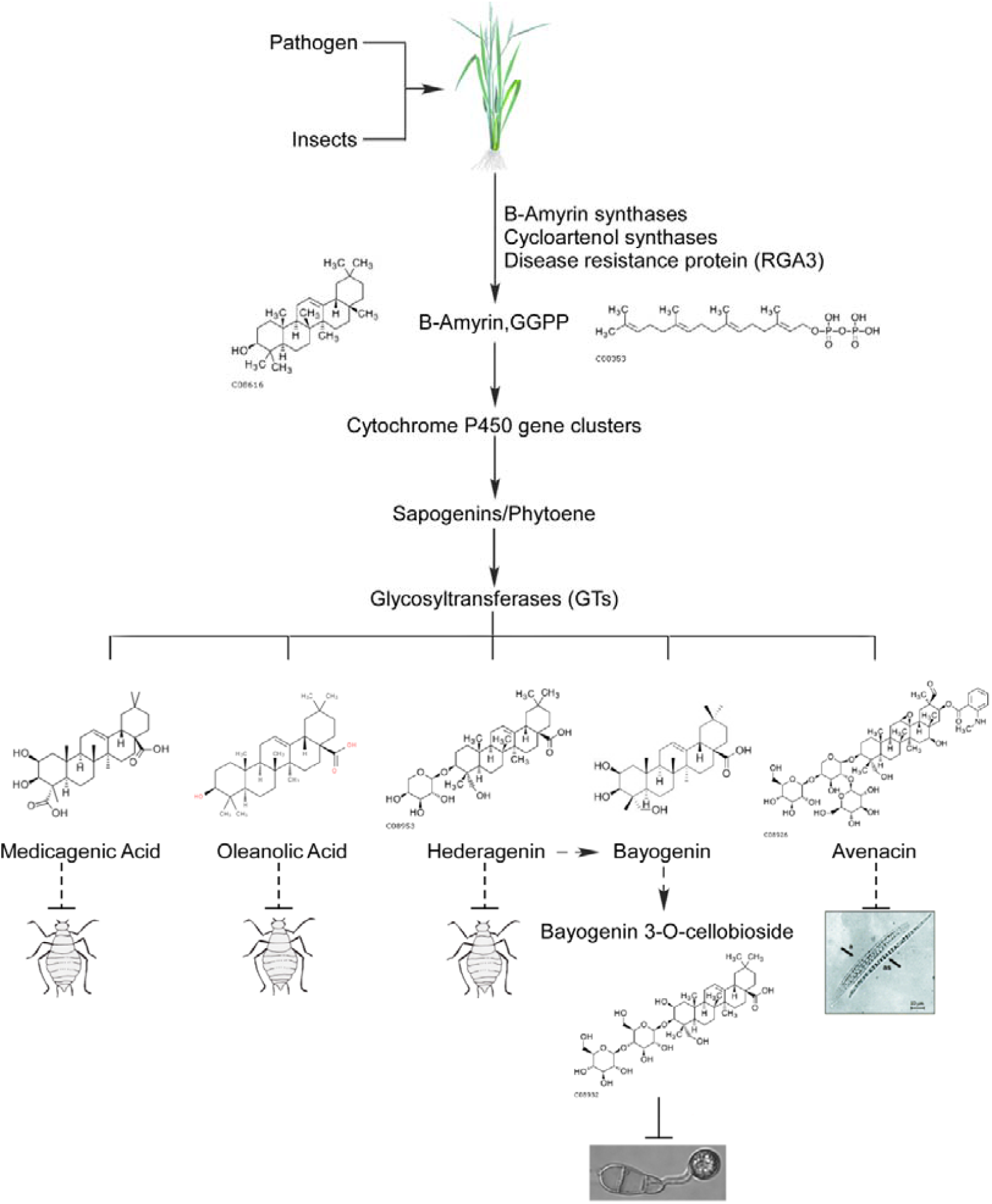
Schematic representation of different types of saponins their exclusive insecticidal and fungicidal effects. *P. oryzae* mediated metabolome-reprogramming result in the generation of Bayogenin 3-O-Cellobioside in resistant and susceptible rice cultivars form both *japonica* and *indica* lineages combined enzymatic activities multiple diphosphate glucuronyltransferases and β-amyrin synthases. Lkie avencin, Bayogenin 3-O-Cellobioside represent a novel rice saponin with anti-blast effect.

## Materials and Methods

### 1. Preparation of rice samples for metabolomics assay

2-3-week-old rice seedlings were sprayed-inoculated with conidia suspensions (1.5-2.0 × 105 conidia/mL containing 0.02% Tween20) and incubated in a humid chamber along with the control groups (sprayed with water containing 0.02% Tween20) for 12 hours at 27 □ Leaf tissues were harvested from respective groups separately (inoculated and non-inoculated groups) 12 hours post-inoculation. The harvested leaf tissues were ground in liquid nitrogen to yield a fine and homogeneously blended powder. 0.1 g of the leaf powder was mixed with 1000 µl of 50% methanol and incubated in shaking incubator for 12 hours at 4□. The contents were centrifuged at 13000 rpm for 15 minutes at 4□. The supernatant was pipetted into new eppendorf tubes and 15 µl of the extracts were diluted 10-fold with 70% (v/v) cold methanol and filtered with 0.2μm Milex Millipore membrane into sample bottles with glass insert. The diluted extracts were stored at 4□ and were later used for non-targeted whole-metabolome analysis. Whole-metabolome profiling data was generated with 2777 C UPLC system (Waters, UK) type of Liquid chromatography and Xevo G2-XS QTOF (Waters, UK) mass spectrometry instruments (BGI.Tech metabolomics platform at ShengZheng). The HPLC assay was conducted with six technical replicates.

### 2. Culturing and preparation of Magnaporthe oryzae mycelia for Metabolomic Assay

Wild-type *P oryzae* Guy11 samples for metabolomics assays were cultured in complete liquid media (CM) (6 g yeast extract, 6 g casamino acid, 10 g sucrose in 1L distilled water) in a shaking incubator operating at 150 rpm at 28.5□ for 5 days. The cultured strains were subsequently filtered and thoroughly rinsed with sterilized double deionized water (ddH2O) and freeze-dried in 70% (v/v) methanol for 24 hours in (Labconco Free Zone 12L). The dried hyphae tissues were ground into powder using a pestle and mortar. 0.16 mg of the ground hyphae was mixed with 1.5mL of 50% (v/v) methanol, vortexed vigorously to yield a uniform mixture, and incubated in a water bath at 65□ for 1 hour. After incubation, the mixture was centrifuged for 10 minutes at 12000 rpm. The supernatant was aliquoted into a new 2.0 mL sterilized eppendorf tube, 15μL of the supernatant was diluted 10-fold with 70% (v/v) methanol and filtered with 0.2μm Milex Millipore membrane into sample bottles with glass insert and stored at 4□ for metabolic analysis.

### 3. Harvesting and preparation of conidia for metabolomic Assay

To generate conidia for metabolomics analysis, a mycelial plug of Wild-type *P* oryzae Guy11 strains were grown on rice bran medium, at 27□ with constant exposure to light. After 10 days the conidia were harvested, washed with sterile distilled water, and was observed under the microscope. The washed conidia were then filtered and centrifuged for 10 minutes at 12000 rpm. The conidia were ground in liquid-nitrogen to yield fine powdered. 0.10 mg of the conidia powder was mixed with 1.5mL of 50% (v/v) methanol, vortexed vigorously to yield a uniform mixture and incubated in a water bath at 65□ for 1 hour. After incubation, the mixture was centrifuged for 10 minutes at 12000 rpm. The supernatant was aliquoted into a new 2.0 mL sterilized Eppendorf tube, 15μL of the supernatant was diluted 10-fold with 70% (v/v) methanol and filtered with 0.2μm Milex Millipore membrane into sample bottles with glass insert and stored at 4□ for metabolic analysis.

### 4. Generation of Appressorium for Metabolomic assays

For appressorium formation metabolome profiling, appressoria were generated by dropping an aliquot of 1.0 mL per of conidia suspension (1×105) on fisher scientific hydrophobic slide surface and incubated in a humid chamber at 26□ without light. Appressorium formation was observed after 12 hours using an optical microscope. Solution drops containing the developed appressorium were pipetted into sterilized EP-tubes and centrifuged for 5 minutes at 5000 rpm. The liquid was pipetted-out and the pellet was transferred, frozen and ground in liquid nitrogen to yield a fine powder using pestle and mortar. 0.10 mg of the powder generated was mixed with 1.5mL of 50% (v/v) methanol, vortexed vigorously to yield a uniform mixture and incubated in a water bath at 65□ for 1 hour. After incubation, the mixture was centrifuged for 10 minutes at 12000 rpm. The supernatant was aliquoted into new 2.0 mL sterilized Eppendorf tubes, 15μL of the supernatant was diluted 10-fold with 70% (v/v) methanol and filtered with 0.2μm Milex Millipore membrane into sample bottles with glass insert and stored at 4□ for metabolic analysis.

### 5. Pathogenicity Assay

For plant infection assays, conidia were collected from strains cultured on rice-bran medium for 7-10 days. Conidial suspensions were adjusted to 1.5-2.0 × 105 conidia/mL in 0.02% Tween solution and sprayed onto 3-4-week-old susceptible (*CO39, LTH* & *NPB*) and resistant (*Pi-b, Pi-4b*, & *Pi-gm*) rice seedlings. Inoculated plants were incubated in a dark, humid chamber at 25□ for 24 hours, and then moved to another humid chamber with a 12 hour photoperiod. The plants were examined for disease symptoms at 7-days post-inoculation.

### 6. Evaluating the influence of rice leaf extracts on conidia germination and appressorium formation

Conidia were collected from 7-day-old rice-bran medium. Conidial suspensions were adjusted to 1.5-2.0 × 105 conidia/mL in 0.02% Tween solution and sprayed onto 3-4-week-old susceptible (CO39, LTH & NPB) and resistant (Pi-b, Pi-4b, & Pi-gm) rice seedlings. Inoculated plants were incubated in a dark, humid chamber at 25□ for 24 hours, then moved to another humid chamber with 12 hour photoperiod. The inoculated rice leaves were then grounded in liquid nitrogen into a fine powder. About 1g of crushed leaves were dissolved in 4 ml of 80% methanol and incubated at 4 °C on a shaking incubator overnight. After overnight shaking, the mixture was centrifuged for 10 minutes at 13000 g to obtain the supernatant. The supernatant was then filtered with non-sterilized millex syringe driven membrane. The substrate syrup was used to directly wash conidia from the culture plates. 20 uL of the conidia suspensions was placed on a fisher scientific hydrophobic microscope cover glass and incubated in a dark humid chamber at 26°C before proceeding to appressorium formation.

### 7. RNA extraction and generation of Illumina RNA sequencing library

Total RNA was extracted from the inoculated rice seedlings (C_Co39, C_NPB, and C_gm) along with their non-inoculated control group T_Co39, T_NPB, and T_gm. The extraction of total RNA from inoculated and non-inoculated control samples was carried-out with RNAprep pure Plant Kit (Tiangen, Beijing) by following processes recommended by the manufacturer. RNA degradation and contamination were measured by running the extracted RNAs on 1% agarose gels. RNA integrity was assessed using the RNA Nano 6000 Assay Kit of the Bioanalyzer 2100 system (Agilent Technologies, CA, USA). The RNA concentration was measured with an RNA Assay Kit in Qubit 2.0 Flurometer (Life Technologies, CA, USA). The RNA concentration was measured with an RNA Assay Kit in Qubit 2.0 Flurometer (Life Technologies, CA, USA)The cDNA library was sequenced on the Illumina sequencing platform (IlluminaHiSeq 2000) with 150 bp pair-end reads length and 300 bp insert size by Gene Denovo Co. (Guangzhou, China). Novogene in-house Perl script was used to select clean reads by removing adaptor sequences, low-quality sequences (reads with more than 50% of bases quality lower than 20) and reads with more than 5% N bases. The reference genome of Nipponbare genome Oryza sativa Japonica and gene model annotation files were downloaded from the genome website directly (58). Index of the reference genome was built using Hisat2 v2.0.4, and paired-end clean reads were aligned to the reference genome using Hisat2 v2.0.4. We selected Hisat2 as the mapping tool for that Hisat2 can generate a database of splice junctions based on the gene model annotation file and thus a better mapping result than other non-splice mapping tools.

## Additional information

Accession codes: Details of the RNA-Seq data generated in this study have been deposited in the NCBI Sequence Read Archive database and can be accessed with the accession code: GSE126961

## Acknowledgements

This work was supported by National Natural Science Foundation of China (No. U1805232, No.31770156) and National Key Research and Development Program of China (2016YFD0300707)

## Notes

https://www.ncbi.nlm.nih.gov/geo/query/acc.cgi?acc=GSE126961

