## Supplementary Table and Figures for "Bayogenin 3-O-Cellobioside is a novel non-cultivar specific anti-blast metabolite produced in rice in response to *Pyricularia oryzae* infection"

### List of Supplemetary Figures and Tables

**Supplementary Table 1:** Resistance characteristics of six different rice cultivars from both *indica* and *japonica* lineage.

| Rice cultivars | Leanage | Resistance level |
|---|---|---|
| CO39 | *Indica* | Completely susceptible |
| NPB | *Japonica* | Completely susceptible |
| LTH | *Japonica* | Completely susceptible |
| Pi-gm | *Indica* | Completely resistant |
| Pi-b | *Indica* | Completely resistant |
| Pi-4b | *Indica* | Moderately resistant |

Table2a. Total metabolites from leaf extract of control and treatment groups of different rice cultivars

| Rice Cultivars | Control | | Treatment | |
|---|---|---|---|---|
| | Negative | Positive | Negative | Positive |
| **CO39** | 13468 | 10437 | 13264 | 10798 |
| **NPB** | 13294 | 10397 | 13298 | 10852 |
| **LTH** | 12919 | 10255 | 12913 | 10808 |
| **Pi-gm** | 13008 | 10325 | 12920 | 10372 |

| | | | | |
|---|---|---|---|---|
| **Pi-b** | 13188 | 10324 | 12967 | 10798 |
| **Pi-4b** | 13295 | 10240 | 13144 | 10864 |

Table2b. Total metabolites recorded in different development stage of *P. oryzae*

| **Guy11** | **Negative** | **Positive** |
|---|---|---|
| **Mycelia** | 1325 | 1524 |
| **Conidia** | 888 | 708 |
| **Appressoria** | 837 | 1799 |

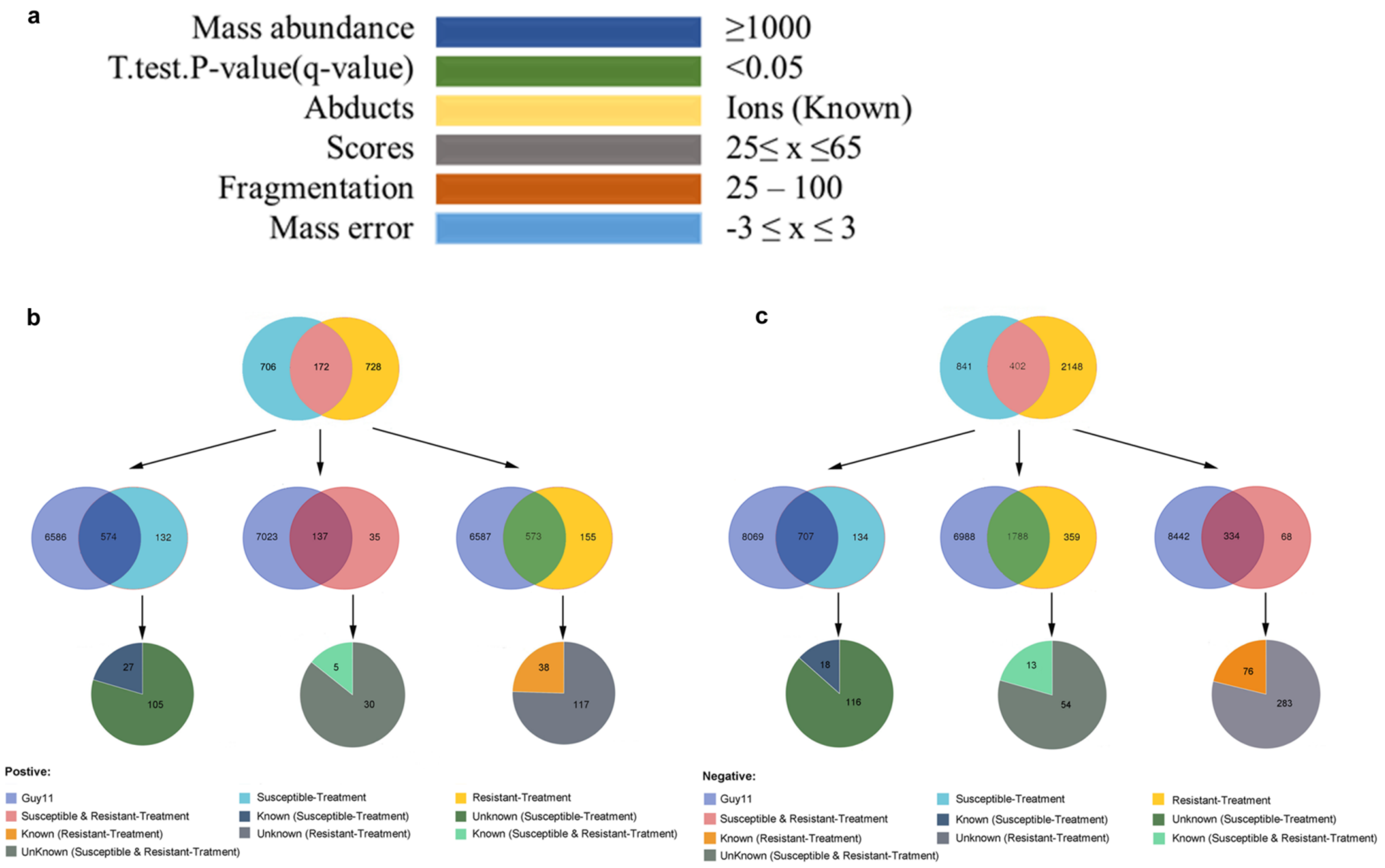
a
Mass abundance ≥1000
T.test.P-value(q-value) <0.05
Abducts Ions (Known)
Scores 25≤ x ≤65
Fragmentation 25 – 100
Mass error -3 ≤ x ≤ 3
b
706 172 728
6586 574 132
7023 137 35
6587 573 155
27 105
5 30
38 117
c
841 402 2148
8069 707 134
6988 1788 359
8442 334 68
18 116
13 54
76 283
Postive:
Guy11
Susceptible & Resistant-Treatment
Known (Resistant-Treatment)
UnKnown (Susceptible & Resistant-Tratment)
Susceptible-Treatment
Known (Susceptible-Treatment)
Unknown (Resistant-Treatment)
Resistant-Treatment
Unknown (Susceptible-Treatment)
Known (Susceptible & Resistant-Tratment)
Negative:
Guy11
Susceptible & Resistant-Treatment
Known (Resistant-Treatment)
UnKnown (Susceptible & Resistant-Tratment)
Susceptible-Treatment
Known (Susceptible-Treatment)
Unknown (Resistant-Treatment)
Resistant-Treatment
Unknown (Susceptible-Treatment)
Known (Susceptible & Resistant-Tratment)

**Supplementary Figure 1**: Different rice cultivars undergo common and differential metabolome reprogramming in response to *P. oryzae* infection. **(a)** Showed the filtering parameters used for eliminating background noises and biologically insignificant metabolites prior to comaparative metabolomics analysis. (b) Showed the metabolomics differentials between non-inoculated rice cultivars, inoculated rice cultivars, and the Guy11 strain under (Pos.$^{+}$) ionization mode. **(c)** Showed the metabolomics differentials between non-inoculated rice cultivars, inoculated rice cultivars, and the Guy11 strain under (Neg.$^{-}$) ionization mode.

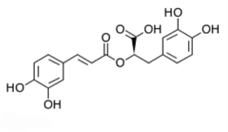

C01850

Rosmarinic acid

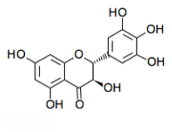

C02906

Dihydromyricetin*

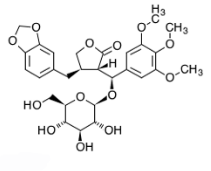

C10876

Podorhizol beta-D-glucoside*

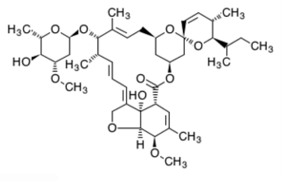

C11982

Avermectin A1a monosaccharide

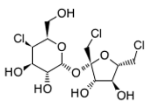

C12285

Sucralose

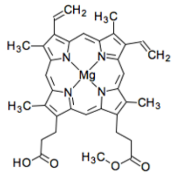

C04536

Mafnesium protorphyrin monomethyl ester

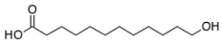

C08317

12-Hydroxydodecanoic acid

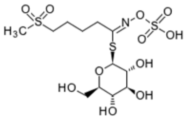

C08410

4-Methylsulfonylbutyl glucosinolate*

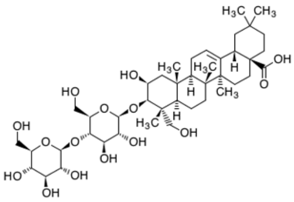

C08932

Bayogenin 3-O-cellobioside*

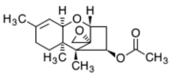

C09741

Trichodermin

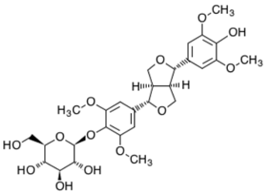

C10890

(+)-Syringaresinol O-beta-D-glucoside*

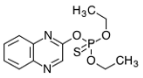

C11030

Quinalphos

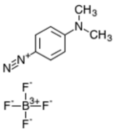

C13681

P-n,n-(Dimethylamino) phenyldiazonium fluoroborate

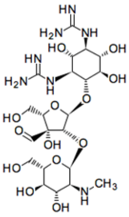

C17571

5'-Hydroxystreptomycin

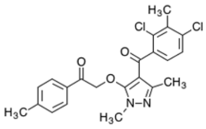

C18555

Benzofenap

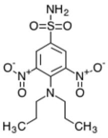

C18877

Oryzalin

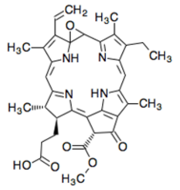

C21192

Epoxypheophorbide A

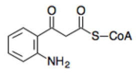

C21452

2-Aminobenzoylacetyl-CoA

**Supplemetary Figure 2: Structure formulae of 19 non-cultivar specific metabolites identified in resistant and susceptible rice cultivars exclusively during *P. oryzae* infection.** Name and KEGG compound identification code for the respective metabolites are written directly under each structural formulae. Compounds with an asterisk (*) and phytochemicals.

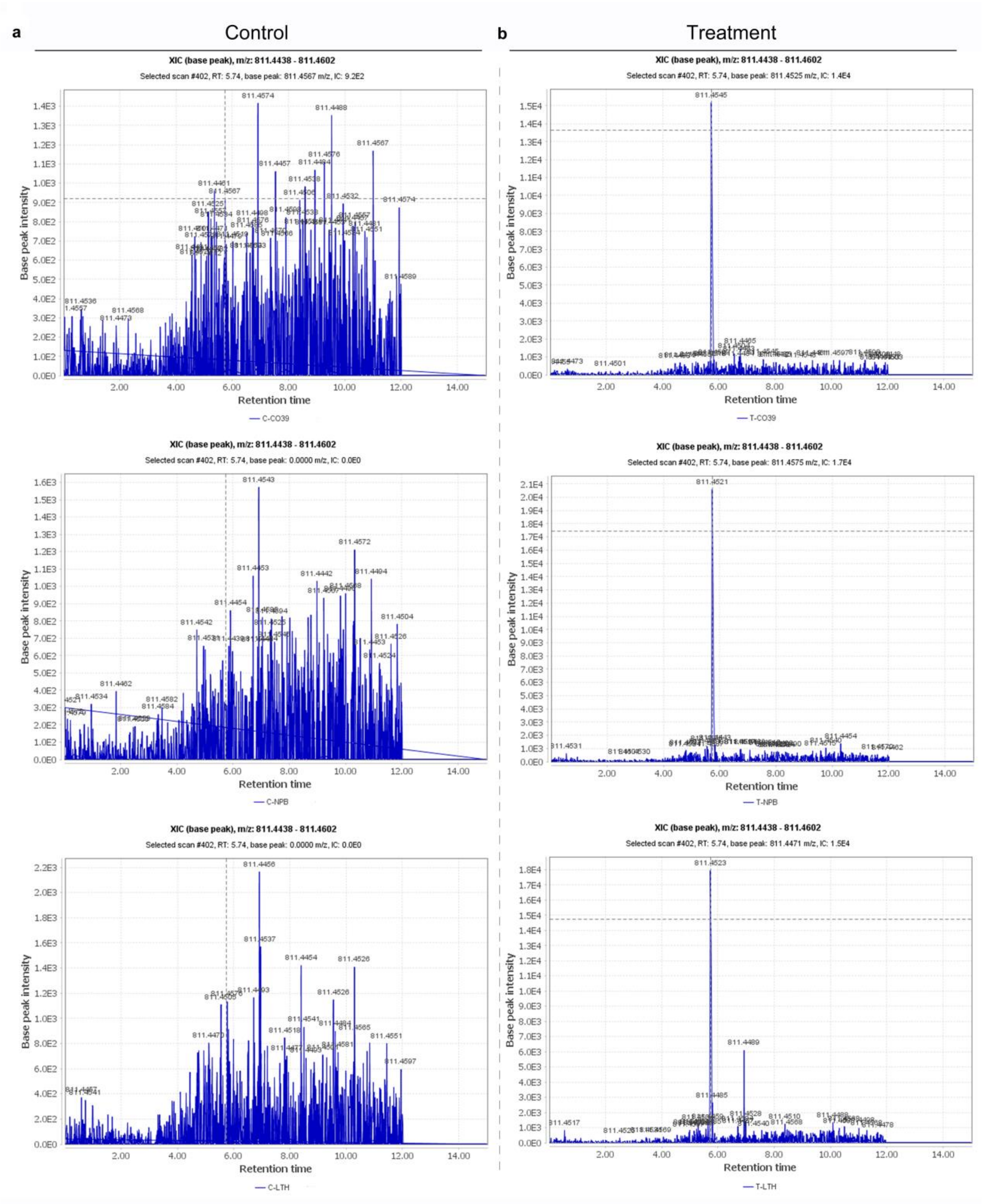
a
Control
b
Treatment
XIC (base peak), m/z: 811.4438 - 811.4602
Selected scan #402, RT: 5.74, base peak: 811.4567 m/z, IC: 9.2E2
Selected scan #402, RT: 5.74, base peak: 811.4525 m/z, IC: 1.4E4
Selected scan #402, RT: 5.74, base peak: 0.0000 m/z, IC: 0.0E0
Selected scan #402, RT: 5.74, base peak: 811.4575 m/z, IC: 1.7E4
Selected scan #402, RT: 5.74, base peak: 811.4471 m/z, IC: 1.5E4
Base peak intensity
Retention time
C-CO39
T-CO39
C-NPB
T-NPB
C-LTH
T-LTH

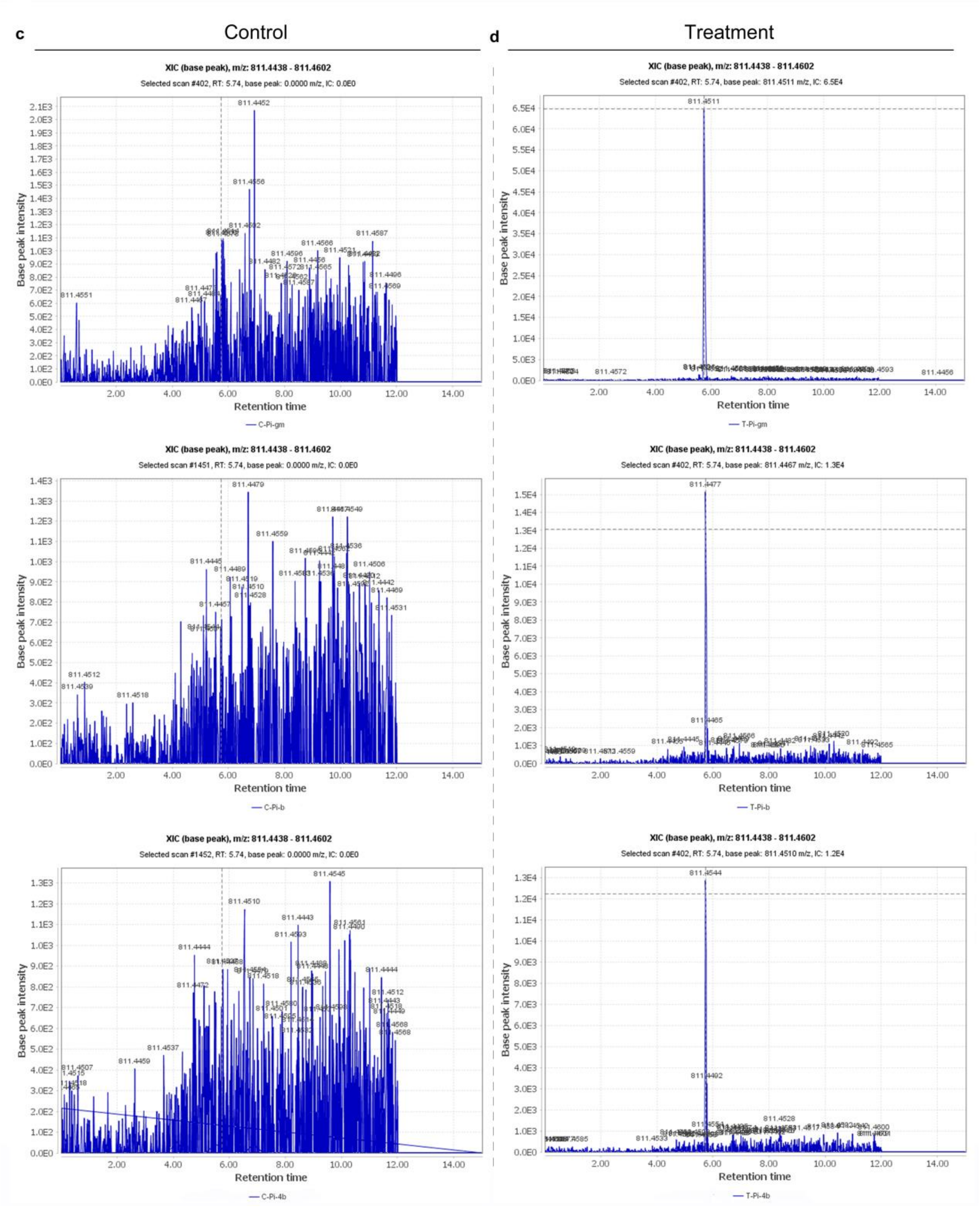


**Supplementary Figure 3: Exclusive generation of Bayogenin 3-O-Cellobioside in P. oryzae inoculated susceptible rice cultivars**. (a) showed the absences of spectra peaks Bayogenin 3-O-Cellobioside in non-inoculated susceptible rice cultivars (CO39, NPB, and LTH) (b) The Mass Spectra ( peak intensity/retention time) showed the detection of Bayogenin 3-O-Cellobioside in susceptible rice cultivars (CO39, NPB, and LTH) in response to P. oryzae infection. (c) Showed the absences of spectra peaks Bayogenin 3-O-Cellobioside in non-inoculated resistant rice cultivars (Pi-gm, Pi-4b, and Pi-b) (d) The Mass Spectra ( peak intensity/retention time) showed the detection of Bayogenin 3-O-Cellobioside in resistant rice cultivars (Pi-gm, Pi-4b, and Pi-b) challenged with P. oryzae. The peaks shown in the chromatogram are those that fall within the 10ppm range of 811.452 within 0-15 min, The base peak represent the target peak. The peaks intensity below the standard ionization response intensity (IC) value is considered as noise signal, and a base peak of “0” means there is no target compound.
